# EXPLORE: A novel deep learning-based analysis method for exploration behaviour in object recognition tests

**DOI:** 10.1101/2022.06.24.497470

**Authors:** Victor Ibañez, Laurens Bohlen, Francesca Manuell, Isabelle Mansuy, Fritjof Helmchen, Anna-Sophia Wahl

## Abstract

Object recognition tests are widely used in neuroscience to assess memory function in rodents. Despite the experimental simplicity of the task, the interpretation of behavioural features that are counted as object exploration can be complicated. Thus, object exploration is often analysed by manual scoring, which is time-consuming and variable across researchers. Current software using tracking points often lacks precision in capturing complex ethological behaviour. Switching or losing tracking points can bias outcome measures. To overcome these limitations we developed ”EXPLORE”, a simple, ready-to use and open source pipeline. EXPLORE consists of a convolutional neural network trained in a supervised manner, that extracts features from images and classifies behaviour of rodents near a presented object. EXPLORE achieves human-level accuracy in identifying and scoring exploration behaviour and outperforms commercial software with higher precision, higher versatility and lower time investment, in particular in complex situations. By labeling the respective training data set, users decide by themselves, which types of animal interactions on objects are in- or excluded, ensuring a precise analysis of exploration behaviour. A set of graphical user interfaces (GUIs) provides a beginning-to-end analysis of object recognition tests, accelerating a fast and reproducible data analysis without the need of expertise in programming or deep learning.

## Introduction

The object recognition test (ORT) has become one of the most frequently used tests in biomedical research to fast and efficiently assess different phases of learning and memory in rodents. Based on an observation by Berlyne [1] in 1950, who described that rats spend more time on a novel object than at a previously explored (familiar) one, Ennaceur and Delacour [2] developed a behavioural task with three phases (habituation, acquisition and testing) to evaluate ”object recognition”: During habituation the animal is exposed to the testing environment, which typically differs from the housing cage, to acclimate to this new environmental condition. During the acquisition phase, the animal is presented and familiarised with a set of objects before turning back to the home cage for a retention period. In the testing phase one of the previously explored, similar objects is replaced by a novel object which deviates in shape from the originally presented (familiar) one.

As rodents have an innate preference for novelty, animals will spend more time exploring the novel object once they have recognized the familiar object(s) [1]. By definition, novelty requires intact memory of the previously experienced object(s). As such, the animal‘s behaviour in the ORT reflects underlying neural processes mediating memory storage and retrieval of features of the familiar object(s) which can be assessed with appropriate recording techniques. The ORT therefore has been applied in various models of aging and different neurological entities with cognitive impairment. Animals were also assessed in the ORT for drug discovery of new pharmacological agents [3–8]. The ORT has several advantages over other rodent memory tasks: As it relies on the natural curiosity for exploring novelty in rodents, neither training of the test nor any positive or negative reinforcement are required. Therefore, the ORT is simpler and less stressful and requires shorter test durations compared to other commonly used memory tests including the Morris water maze or the Barnes maze [8]. Furthermore, the task can be repeatedly performed with the same animal under the same or modified conditions assessing the acquisition, consolidation and retrieval of memory.

However, despite the simplicity of the experimental implementation of the test, the analysis is rather time-consuming and has been liable to subjectivity: Traditionally, the scoring of the time the animal spends exploring each object, is done manually - either during the experiment or post-hoc based on video recordings. In addition, which type of interactions are part of “exploring behaviour” and which are not, is a matter of debate. For example, when the animal turns its nose towards the object, up to which distance from the object does this sniffing behaviour count as exploration? As a result, manual scoring of the behaviours of interest may vary across researchers, especially if new researchers are trained [9]. Even scoring of multiple videos and sessions by the same investigator can be subject to bias.

To overcome these limitations and increase the reliability of results, it would be a significant advance if a researcher could define a set of “explorative behaviour patterns” and then use software to identify how frequently these behaviours occurred at a given object in a video. Early video tracking programs [10] tracked only the center point of an animal, compromising the reliability of the analysis of the ORT, during which multiple body points are involved in object exploration and thus should be tracked simultaneously. Besides costly, commercial software (EthoVision, Viewpoint, ANY-maze) perform tracking of multiple body points with a focus on the nose-point detection capabilities [10, 11], enabling scoring of distinct forms of “sniffing” behaviour. Drawbacks of these methods are that other interactions with the objects, such as climbing or hiding under the object, may or may not be counted as “exploring behaviour”. If only nose detection within a certain radius surrounding an object is considered as exploration, a significant amount of “exploring behaviour” may be missed.

Within the past decade, progress in the field of machine learning – especially deep learning – gave rise to new methods for image analysis: Instead of using tracking-points, convolutional neural networks extract important features of given objects in images. The task of automatically classifying the action of an animal into user-defined behaviours falls into the category of supervised machine learning.

Newer solutions including JAABA [12], SimBA [13] and others [14] have further improved the capabilities of supervised classification of behaviour, using the time series of specific behavioural features to classify whether a behaviour is present at a given time point, with further refinement including pose estimation methods [15–17] or purely pixel based approaches [18]. Although these recent methods are powerful in detecting and quantifying behaviour, they typically are designed to analyse motor behaviour and are not developed for the specific needs of analyzing object exploration in the ORT. In particular, they lack a user-friendly stopwatch function to score the time duration that animals spend exploring each object, which is a prerequisite to calculate the discrimination index used by many labs to quantify and compare memory function [19, 20].

As a solution to this problem, we here present EXPLORE, a software tool which was specifically developed and designed for the analysis of explorative behaviour in open fields or in the ORT. EXPLORE is a deep-learning based tool, which allows users to decide by themselves which object interactions of the animal is accounted as ”explorative behaviour” by labeling the appropriate training data. Based on these training data a convolutional neural network then extracts features from images and classifies them into the predefined classes. Exploration is classified with above 95% accuracy on average over many experimental videos, matching expert-level human performance. The robustness of our approach is validated through testing on seven data sets, with training data labeled from different experts, on mouse strains with different colours and recordings in different light and environmental conditions. High performance is achieved by labeling only a few frames of training data and even complex ethological behaviours (e.g. stretching the body or sitting on an object) can be captured. Importantly, specialized video recording hardware is not required, and the entire pipeline does not depend on programming skills by the end-user. We introduce a set of graphical user interfaces (GUIs) for annotating videos, training models, and generating predictions, which can be found on GitHub. Our method also requires relatively low computational resources in case researchers do not have access to large computing clusters or high-end GPUs.

## Results

While the object recognition test is a simple task to perform, there are numerous uncertainties when preparing and analysing the test, such as what objects to use and how to define exploration behaviour (see Fig. 1A for a typical analysis setup). Automated analysis software often lack the precision to capture diverse behavioral features, which experimenters have defined for their analysis, potentially leading to erroneous results. Thus, researchers often rely on manual scoring, which in turn is time-consuming and variable across investigators. To overcome this hurdles, we developed EXPLORE, an easy-to-use pipeline making use of supervised deep learning, combining the advantages of both: manual scoring of user defined exploration behavior and consecutive automation. We first evaluated the human performance and inter- and intra-variability between different raters. We then compared the performance of EXPLORE with two standard software used in many neuroscience labs (EthoVision and ANY-maze) and assessed its versatility and possible problems of misclassifications.

**Fig 1.**
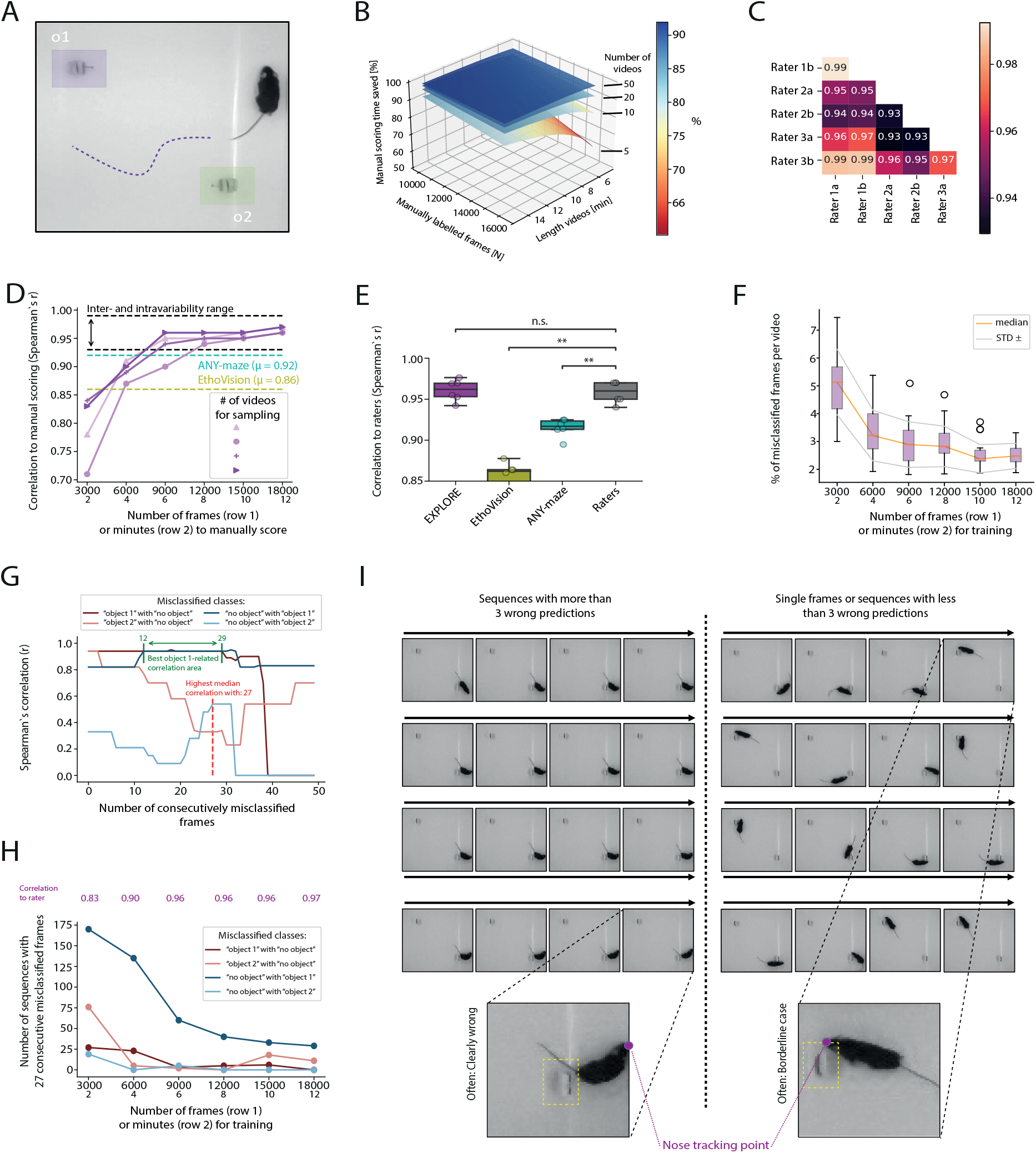
Performance of EXPLORE and misclassifications. **A:** Results are shown for a standard experiment with 2 objects in a quadrangular arena. **B:** 3D-plot showing the % of time saved using EXPLORE as a function of manually labeled frames for training and the length of the experimental videos (coloured planes represent the amount of videos in the experiment): As the amount of videos increases, the time saved also drastically increases.) **C:** Heatmap showing the correlation of each human rater to the other raters. **D:** Line plot showing the mean correlation of EXPLORE to the raters when using different amounts of videos (purple lines) and selecting different amounts of frames (x values) for training data: EXPLORE converts into the human intra- and inter variability range after only a few frames while commercial software performs worse (correlation of ANY-maze is shown in cyan, and for EthoVision in yellow). **E:** Boxplot showing the difference in precision for each tool (paired wilcoxon test with bonferroni post-hoc): Results obtained from EthoVision and ANY-maze were significantly different from human expert rating while no significant difference was found for the automatic quantification of exploration behaviour by EXPLORE. Asterisks showing significance level: ns *p* ≥ 0.05, ** *p* < 0.01. **F:** Boxplots showing the decrease of % of misclassified frames per video as a function of training data labeled (data taken from 15 videos). **G:** Correlation of the accuracy of EXPLORE to human raters as a function of number of consecutively misclassified frames, shown for each of the types of misclassification that occured in the experiment (data taken from 15 videos): “object 1”-related misclassifications (confused classes “object 1” with “no object” and vice versa) with sequences from 12 to 29 frames correlated best. The median of all assessed types of misclassification correlated best at sequences with a length of 27 frames. **H:** Absolute numbers of cases with 27 consecutively misclassified frames in the experiment as a function of training data labeled (data taken from 15 videos): The reduction of longer sequences with (here 27) consecutively wrong predicted frames drives accuracy (see accuracy as correlation to rater in magenta). **I:** Left: Longer sequences of consecutively misclassified frames are often clearly wrong. Right: Shorter sequences of consecutively misclassified frames or single misclassified frames are often boarderline cases (data taken from 15 videos).

### EXPLORE achieves expert-level human performance with little training data while saving research time

Manual scoring is always subject to inter-and intra-rater variance. Thus, we analysed the inter- and intra-rater variability between 6 manual scoring sessions of experienced raters (3 raters, 2 sessions each). Fig. 1C shows that the range of both, inter- and intra-rater variability is between *r* = 0.93 and *r* = 0.99. We argue that every method that aims to compete with human accuracy should at least reach an accuracy within this range. Next, we asked how many frames of manual labeling from how many experimental videos are required to train the network to human accuracy inference on the total (unseen) amount of experimental videos. To this end, we trained EXPLORE a total of 24 times with sampling from 2, 5, 10 and 15 videos (from a data set of 15 videos with 25 fps) by using between 3000 to 18000 manually labeled frames (2 - 12 min) (Fig. 1D). When training with at least 12000 frames (8 min), EXPLORE reached human level accuracy in identifying object exploration behaviour regardless of the number of videos used for sampling. In 3 out of 4 cases (2, 10, 15 videos), EXPLORE reached human accuracy already after 9000 frames (6 min). The variance between videos for sampling stabilized at around *r* = 0.95 after 12000 frames (8 min).

Another important aspect of any automated approach is the time needed to achieve reliable results comparable to human performance. While manual scoring is still the most safe and precise way to analyse object recognition tests, it is very time-consuming. The total time saved when using EXPLORE in comparison to manually scoring all experiment videos is a function dependent on the amount of frames to manually score for training, the length of the experiment videos and the number of videos in the experiment. For example: when manually labeling 15000 frames (10 min) from an experiment with 20 videos with a length of 6 minutes each, a researcher saves around 92% of time compared to manually scoring all of the videos (see Fig. 1B). Assuming it would take the researcher an average of 15 minutes to label a video, then it would take 6 hours of manual scoring in total, while with EXPLORE it would take around 25 minutes. Training our network and applying it to the unlabeled rest of the data depends on hardware and software components of the computer used as well as the steps needed for the model to converge to a global or local minimum. When training with a GPU on Windows the training and prediction took a maximum of 10 minutes each (for 15 videos). With a CPU on MacOS the training took on average 2.5 hours. Training and prediction with EXPLORE required approximately the same time as experienced with commercial software for automated analysis of object recognition tests (EthoVision and ANY-maze).

### EXPLORE outperforms commercial software in correctly detecting object exploration behaviour

Next, we assessed if results scoring object exploration behaviour with EXPLORE correlate to human expert-level and to the most commonly used commercial analysis software (ANY-maze, EthoVision, Fig. 1D). EXPLORE outperformed commercial software (ANY-maze and EthoVision) in a standard experiment as performed by many neuroscience labs (using C57BL/6 wildtype mice, a quadrangular, white arena illuminated with red LEDs to reduce the stress level of the nocturnal animals) with Spearman‘s correlation to human expert scoring of *r* = 0.92 *±* 0.01 and *r* = 0.86 *±* 0.01, respectively. EXPLORE performed clearly within the human accuracy range by achieving an average correlation of *r* = 0.96 *±* 0.01 (see Fig. 1B). We compared the different correlation coefficients by calculating paired Wilcoxon tests, where only EXPLORE showed no significant difference to the human rater values (ANY-maze, EthoVision: *p* < 0.01, see Fig. 1E).

### The accuracy of EXPLORE can be substantially improved when disposing wrongly predicted longer sequences

As results from deep learning methods are sometimes hard to interpret, we analysed one of the experiments from Fig. 1D (random sampling from 15 videos) in more detail. In this experiment there are three classes, which the network can predict for any frame given: ”object 1” (exploration on object 1), ”object 2” (exploration on object 2) and ”no object” (when the rodent is not exploring on either of the objects). Thus, there are six possible scenarios of how the network can mistake the real class (scored by a human rater) for the predicted class (prediction of the network) at a given frame. We refer to these scenarios here with the term ”misclassifications”: (1) The network predicts the class ”object 1” although the real class is ”object 2” or vise versa. (2) The network misclassifes ”object 1” or ”object 2” although it should be ”no object”. (3) ”no object” is classified although the network should have detected either ”object 1” or ”object 2”.

We investigated how these misclassifications influence the accuracy of EXPLORE (Spearman’s correlation to rater) when using more and more training data (taking 3000 to 18000 frames in steps of 3000). First, we extracted the misclassified frames and calculated the percentage of misclassified frames per video for each of the amounts of data taken for training (Fig. 1F). Naturally, one would expect a higher accuracy when less frames are misclassified: we found a strong drop of the median of all videos from 3000 frames (2 min) to 6000 frames (4 min), which explains the major rise of accuracy (Spearman’s correlation to rater) from *r* = 0.83 at 3000 frames of training data to *r* = 0.90 at 6000 frames used for training. However, the cause for the further improvement of the accuracy level from *r* = 0.90 (Spearman’s correlation) at 6000 frames to *r* = 0.96 at 9000 frames remained elusive. Thus, we examined the different possible types of ”misclassifications” (1-3) as described above. We found that errors only occurred at the object locations: the network precisely discriminated between the classes ”object 1” and ”object 2” suggesting a strong robustness for EXPLORE to predict the right location. We then concentrated on the remaining four possible misclassifications (”object 1” with ”no object”, ”no object” with ”object 1”, ”object 2” with ”no object” and ”no object” with ”object 2”). While visually inspecting the misclassified frames, we found two possible types of prediction errors to distinguish: Prediction errors of exploration behaviour which are obvious for every human rater and subtle errors (”borderline cases”), where objective features to classify the frame as object exploration behavior or not were missing (see Fig. 1I). While the first type unarguably compromises the results, the second type would not substantially influence results, since there is no consensus among raters about the frame being either of the two possible classes.

We then quantitatively assessed whether longer consecutively misclassified sequences (*>* 3 frames) or shorter sequences (*<*= 3 frames) and single frames are causal for the low accuracy. Therefore, we calculated the Spearman’s correlation between the accuracy (Spearman’s correlation of EXPLORE to raters) and different numbers of consecutively misclassified frames (from length 0 to 50) for each misclassification (see Fig. 1G). We found that ”object 1”-related misclassifications (”object 1” with ”no object” and vice versa) and in particular sequences of 12 to 29 frames correlated best to the accuracy scores. When calculating the median for all of the assessed misclassifications, sequences of consecutively misclassified frames with length 27 correlated best to the accuracy scores. We then investigated the development of the misclassifications with a sequence length of 27 frames when including more training data. We found a substantial improvement of the accuracy of EXPLORE when reducing ”object 1”-related misclassifications which appeared in longer, consecutive sequences of frames (see Fig. 1H).

### EXPLORE is able to well predict exploration behaviour even under difficult light conditions

A strong contrast between experimental arena, animals and objects is a prerequisite for the success of analysis software, such as EthoVision and ANY-maze, which is based on tracking different points of the animal. Using commercial software these conditions must be taken into consideration when already preparing the object recognition task.

Thus, we examined how well EXPLORE performs under extreme light conditions (low contrast): We used a dataset with a white mouse in a white arena (setting 1) as well as a black mouse in a black arena (setting 2, Fig. 2A). EthoVision and ANY-maze were both not able to process this data. The first inference of EXPLORE on 12000 labeled frames was also rather poor (*r* = 0.6 for setting 1, *r* = 0.58 for setting 2)(Fig. 2B). However, after the first correction step (increasing the amount of labeled training data for the two objects, 2077 frames for setting 1 and 1360 frames for setting 2, respectively), the correlation to the human rater increased to 0.69 in setting 1, whereas the correlation to the human rater decreased to 0.21 in setting 2 (Fig. 2C). After the second correction step, adding more labeled training data (1700 frames for setting 1 and 6078 frames of setting 2), showing no object, the correlation to the human rater increased to 0.81 in setting 1 and to 0.77 in setting 2 (Fig. 2D). When tracking outliers (Fig. 2, red labeled values), we also found an improved correlation for these outliers during the course of correction.

**Fig 2.**
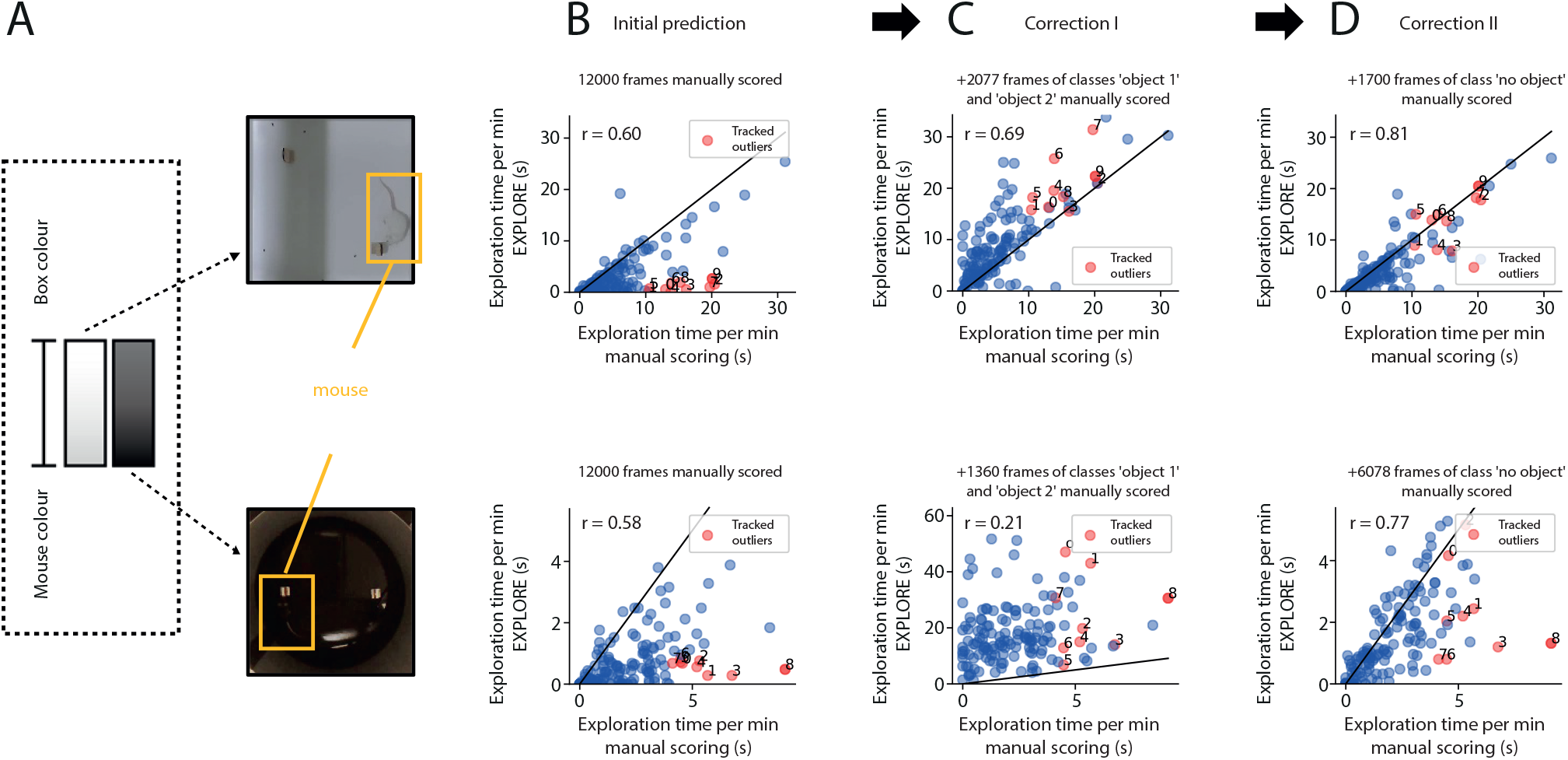
EXPLORE is able to perform well even under difficult light conditions. **A:** Results are shown for a standard experiment with 2 objects in a white quadrangular arena with white mice and in a black round arena with black mice. **B:** Scatterplots indicating the performance of EXPLORE for the initial prediction with 12000 labeled frames for training and its correlation to the human rater for each of the light settings. Outliers are labeled in red and numbered to track them during correction steps. **C:** Scatterplots of the prediction after adding training data with frames specifically labeling the objects of the arena (”object 1” and ”object 2”). **D:** Scatterplots revealing the prediction performance of EXPLORE after adding another training data set without any object (”no object”) and its correlation to the rater after re-prediction. In particular the correlation of outliers (in red) strongly improved compared to B.

### EXPLORE can detect exploration behaviour on a wide range of objects and behavioural features

Usually the ORT is performed with two objects. However, depending on the research question, more objects may be added to the experimental arena. Here we assessed how EXPLORE performs on data with more than two objects (Fig. 3A). Overall EXPLORE performed well with a correlation score of *r* = 0.95 to the human rater. We found very moderate variability when assessing Spearman‘s correlation for each object (”object 1”: *r* = 0.97, ”object 2”: *r* = 0.87, ”object 3”: *r* = 0.98)(Fig. 3B).

**Fig 3.**
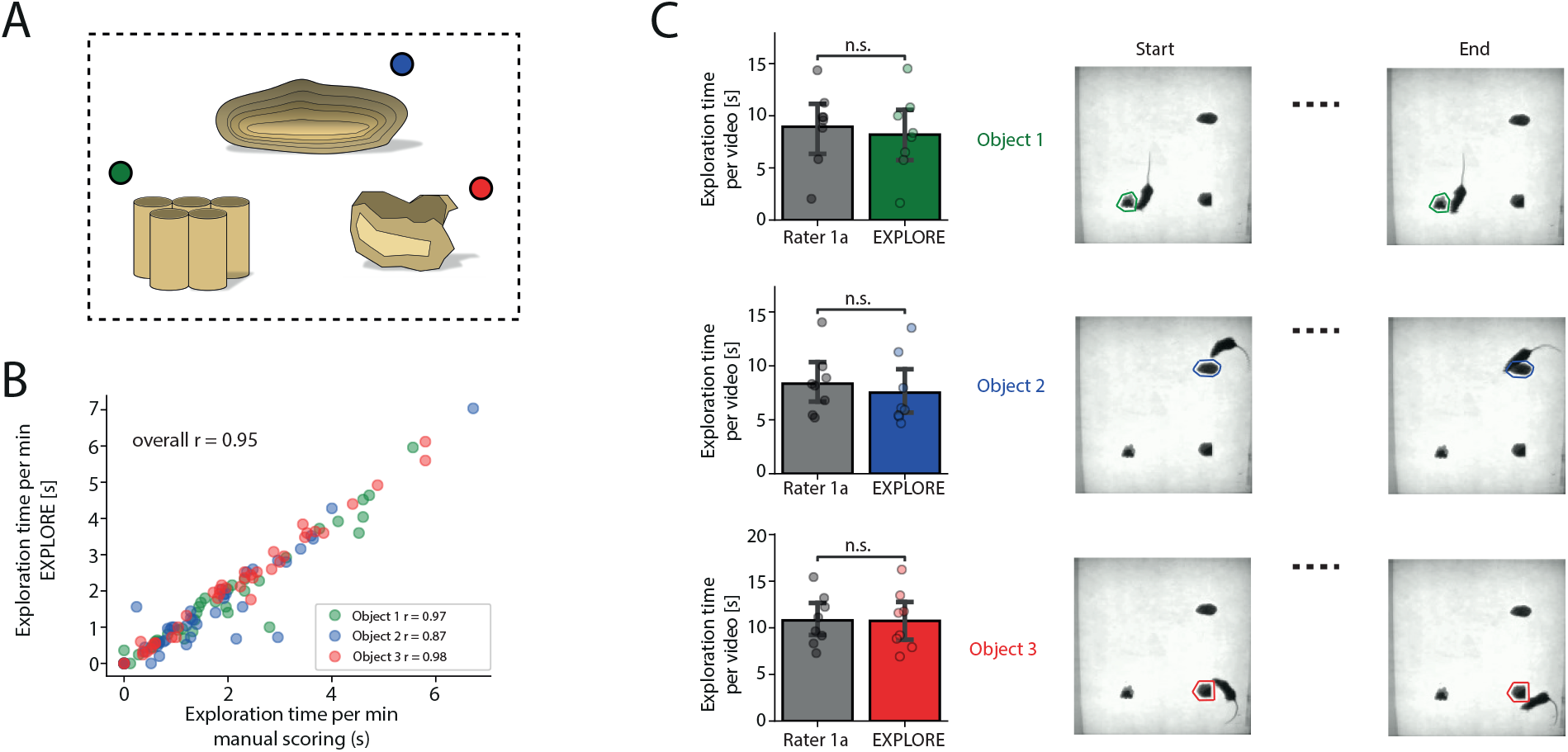
EXPLOREs performance on multiple objects. **A:** Results are shown for an experiment with 3 objects to explore in the experimental arena. **B:** Scatterplot showing how the results of EXPLORE calculating the object exploration time correlates to the human rater level for all objects. **C:** For each object, the exploration time per video measured by EXPLORE was compared to the manually scored exploration time. We found no statistically significant difference between EXPLORE and the human rater (paired Wilcoxon test, the significance level was set to *p* ≥ 0.05 as non significant (n.s.)).

The overall time an animal spent exploring is important, since when too short it might get excluded from the experiment. A high correlation does not necessarily mean that the assessed exploration times also have the same length, but rather that the proportions are similar. For example: *L*_1_ = [1,2,3,4,5] and *L*_2_ = [10,20,30,40,50] have a Spearman‘s correlation coefficient of *r* = 1, but the total length (sum) of *L*_1_ is 15 and the one of *L*_2_ is 150. When comparing the total exploration time as measured by EXPLORE to the one manually scored, we found no statistically significant difference (paired Wilcoxon test, Fig. 3C).

While software based on tracking points often relies on pure geometrics (e.g. calculating the distance between the nose tracking point and the object), the users are unable to define by themselves, which type of behaviour they would like to in- or exclude as object exploration. EXPLORE allows this feature: We show that EXPLORE correctly classifies animals sniffing at one of the objects (classes ”object 1” and ”object 2”), or animals stretching their bodies at one of the objects (class ”stretching”) as well as animals sitting on one of the objects (class ”on object”)(Fig. 4A). EXPLORE achieved an overall correlation to the human rater of 0.95 with negligible variability concerning the different correlation rates for each behaviour (”object 1”: *r* = 0.83, ”object 2”: *r* = 0.89, ”stretching”: *r* = 0.9, ”on object”: *r* = 0.97)(Fig. 4B). We also examined again if EXPLORE quantified the exploration time in the range of a human rater: For all three behavioural features (sniffing on objects, stretching and sitting on the object) we did not find a significant difference between the time measured with EXPLORE and manual scoring (paired Wilcoxon test, Fig. 4C).

**Fig 4.**
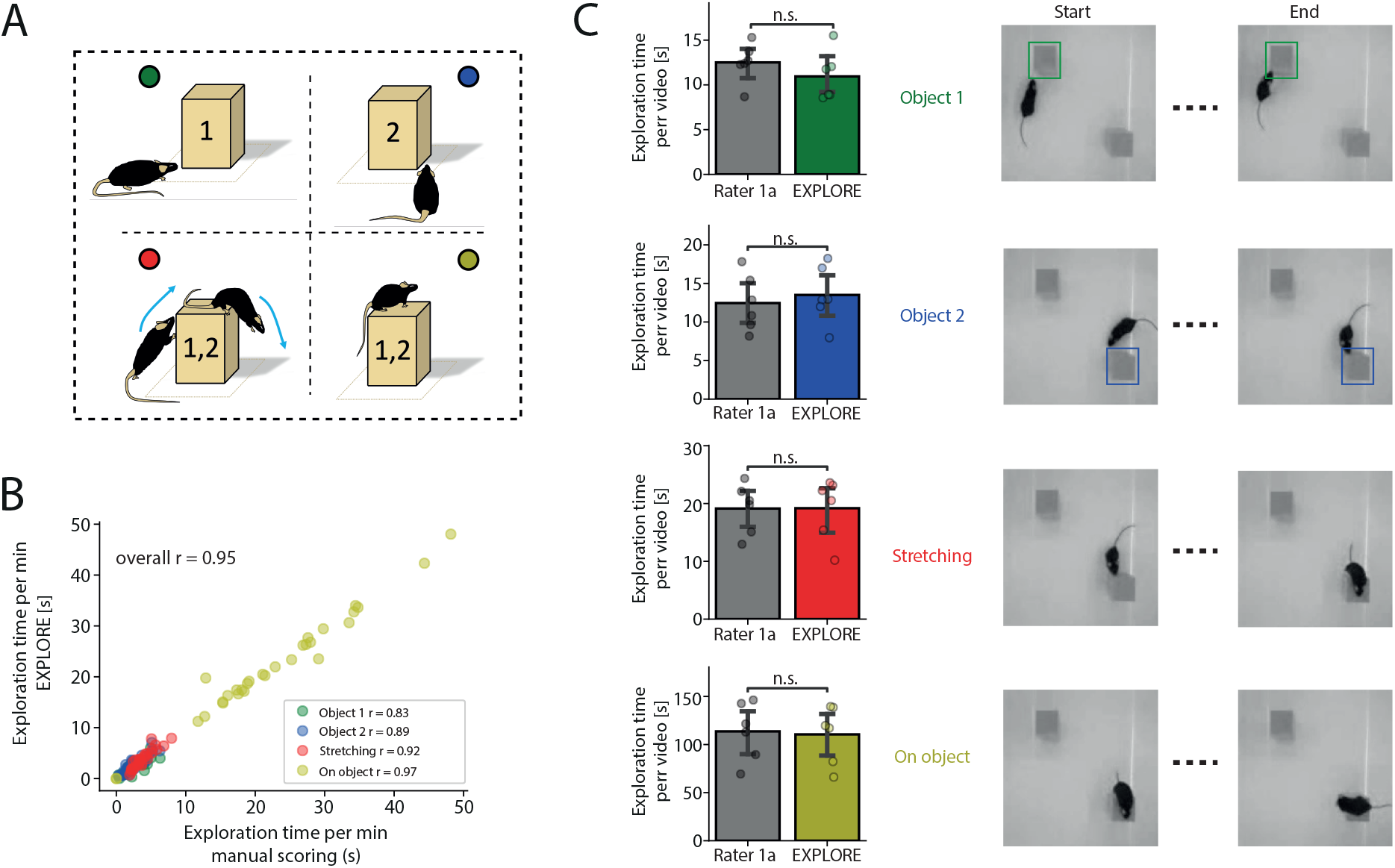
EXPLORE can capture complex behaviour. **A:** Performance of EXPLORE is shown to detect and quantify specific behaviour such as sniffing on objects (1 or 2), stretching the body at the object and sitting on the object. **B:** Scatterplot showing results achieved with EXPLORE correlate to human manual scoring for each type of behaviour. **C:** No significant difference was found between EXPLORE and manual scoring after comparing the time quantified for each specific behaviour (paired Wilcoxon test, the significance level was set to *p* ≥ 0.05 as non significant (n.s.)).

### EXPLORE performs well in situations where keypoint tracking software fails

Keypoint tracking software is commonly used in neuroscience labs to quantify object exploration behaviour. However, this kind of software is vulnerable to typical perturbations of the experimental set-up, which we are assessing in the following and which may compromise the results of the ORT: (1) Animals are often reflected by the walls of the arena causing a loss of tracking points on the real rodent to the mirror image on the wall (Fig. 5A). (2) Using objects with similar colours to the fur of the animal may lead to switching of head- and tail tracking points when animals are climbing at these objects (Fig. 6A). (3) Sitting and resting on the object may be included as exploration behavior contorting the ”true” exploration time of the object. We show in particular that the ratio between body size of the animal and object size influences the accuracy of detecting exploration behaviour (Fig. 7A). For all of the aforementioned situations we assessed the performance of EXPLORE - in comparison to a human rater and commonly used keypoint tracking software (EthoVision and ANY-maze).

**Fig 5.**
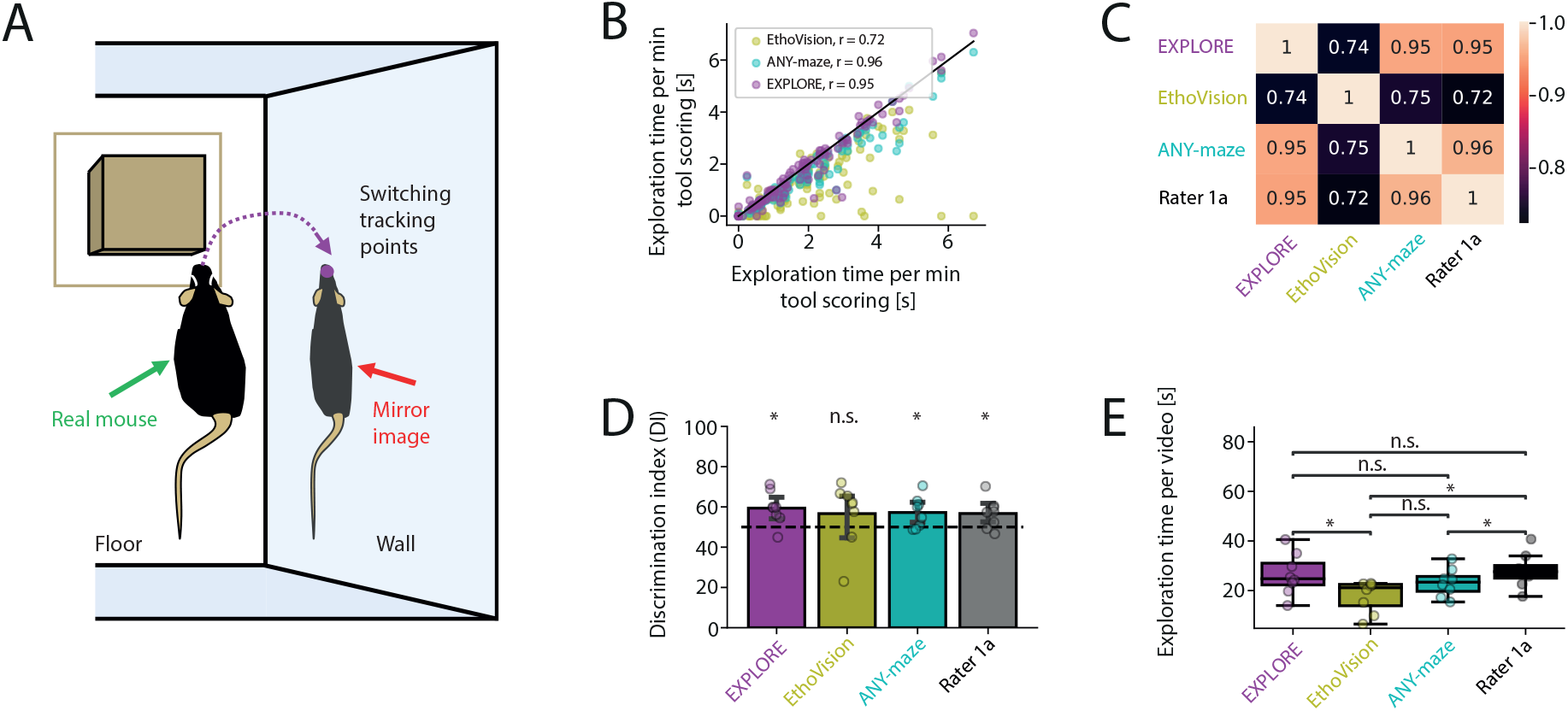
Tracking point loss due to reflecting walls of the experimental set-up. **A:** Problem scheme of software using tracking points: Keypoint tracking of the head of an animal switches to the reflected image of the animal on the wall compromising data analysis. **B:** Scatterplot showing the Spearman‘s correlations of exploration times measured by EXPLORE or the commercial software EthoVision and ANY-maze with human manual scoring. **C:** Heatmap depicting an overview of Spearman‘s correlations between the assessed methods. **D:** Calculation of the Discrimination indexes (DIs) for each of the methods including the human rater and testing against the 50%-chance level: EthoVision provides results not significantly different from the 50%-chance level (Wilcoxon test with Bonferroni post-hoc). **E:** Boxplots of measured object exploration time per video for each method as well as the human rater: Only EXPLORE is not significantly different from the rater (paired Wilcoxon tests were used. Asterisks showing the level of significance: n.s. *p* ≥ 0.05, * *p* < 0.05).

**Fig 6.**
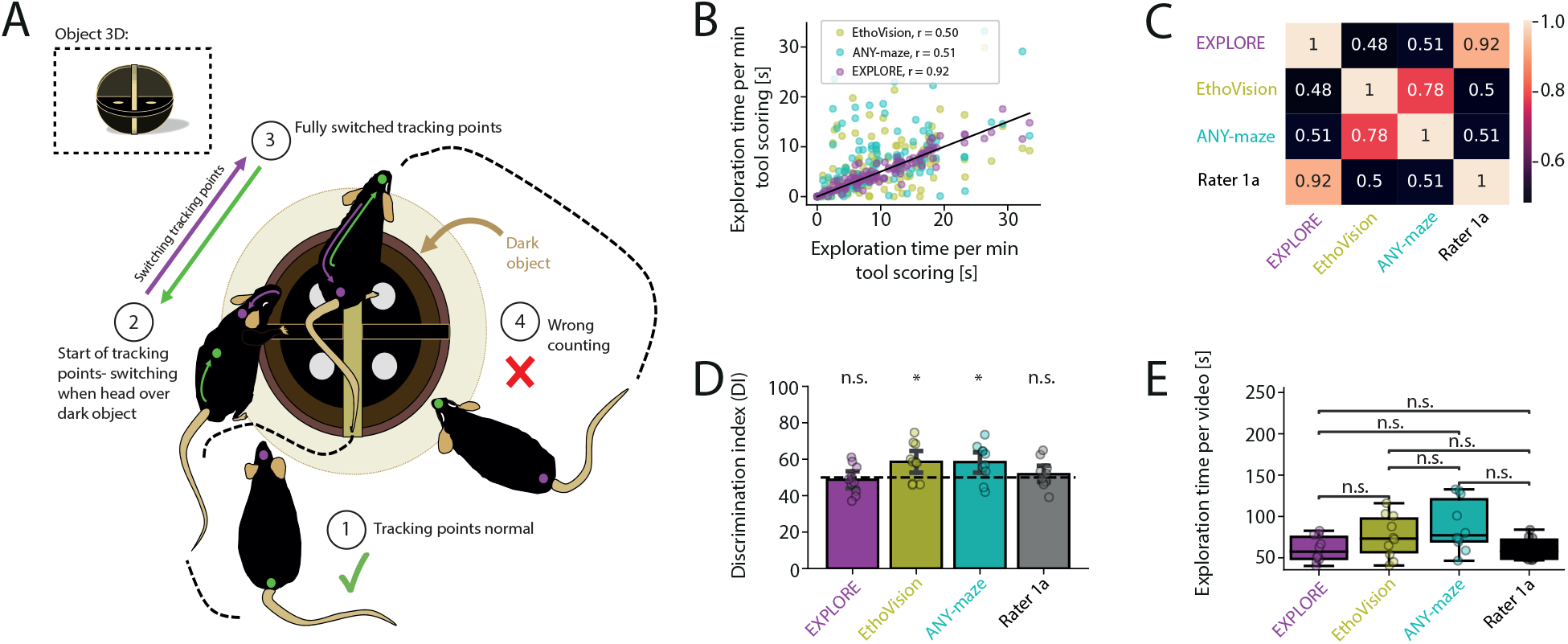
Switching of head- and tail tracking-points when rodents have a similar colour to the object they explore. **A:** Problem scheme showing switching head- and tail tracking-points. **B:** Scatterplot depicting the Spearman‘s correlations of exploration times measured by EXPLORE or the commercial software EthoVision and ANY-maze with human manual scoring. **C:** Heatmap revealing the Spearman‘s correlations between the assessed methods. **D:** Discrimination indexes (DIs) calculated for each of the methods as well as the human rater and tested against the 50%-chance level: Tracking point loss results in a significantly different result for ANY-maze and EthoVision (paired Wilcoxon tests). **E:** Testing boxplots of exploration time per video for each method as well as the human rater against each other: We found no significant difference from the human rater for all methods (paired Wilcoxon tests were used, asterisks showing the level of significance: n.s. *p* ≥ 0.05, * *p* < 0.05.)

**Fig 7.**
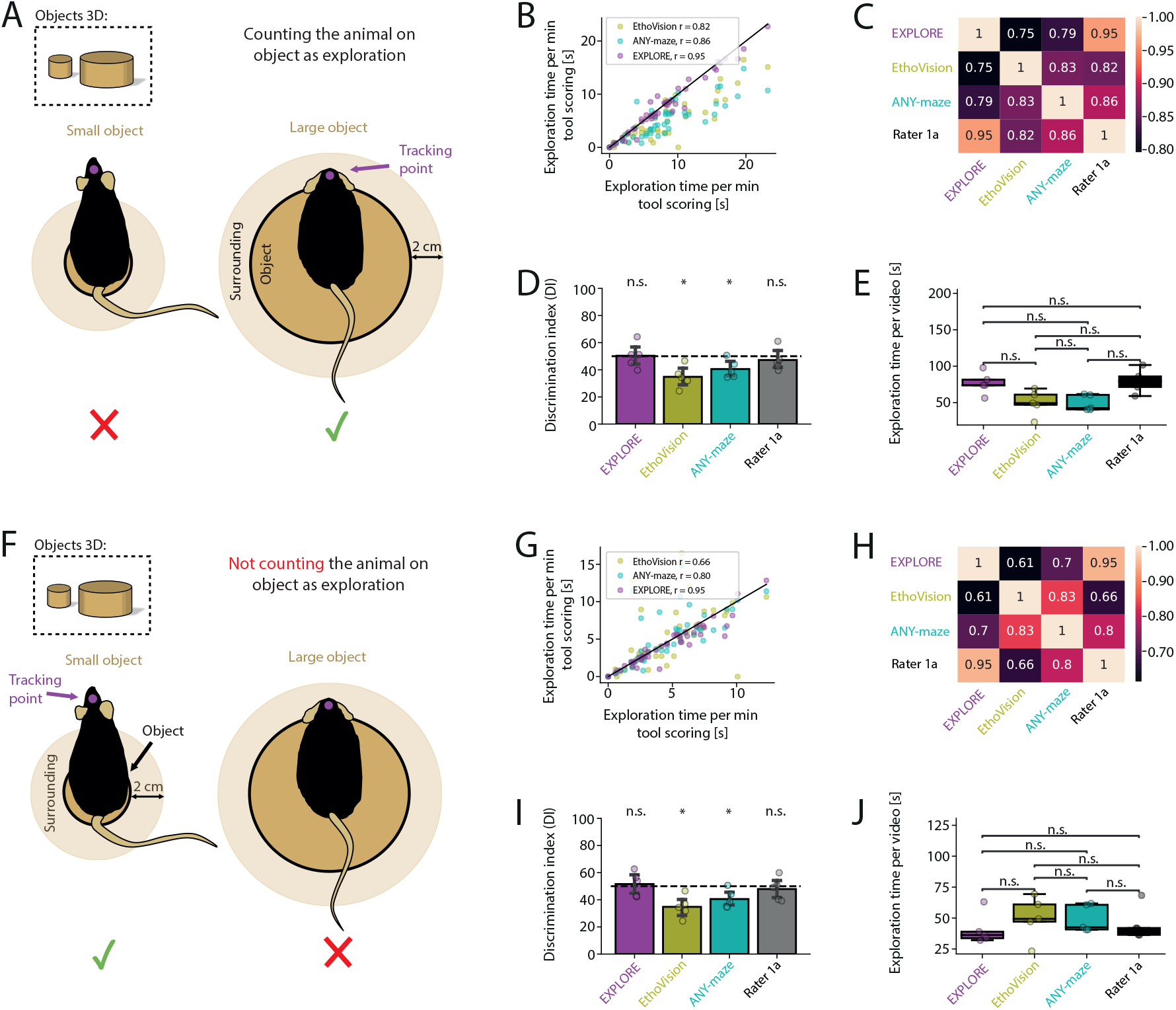
The ratio between object size and body length of the animal influences results of object exploration. **A:** Problem scheme how small and large objects can compromise results of object exploration. Left: Tracking points of an animal sitting on a small object are excluded for calculating the exploration time, as the head of the animal is outside the contour of the object. Right: Tracking points of an animal resting on a large object may be included by commonly used keypoint tracking software. **B:** Scatterplot showing the Spearman‘s correlations of exploration times measured by EXPLORE and the commercial software EthoVision and ANY-maze with human manual scoring. **C:** Heatmap revealing the Spearman‘s correlations between the assessed methods. **D:** Discrimination indexes (DIs) calculated for each of the methods as well as the human rater and tested against the 50%-chance level: Tracking point loss results in a significantly different result for ANY-maze and EthoVision (paired Wilcoxon tests), which is significantly different from the range of EXPLORE and the human rater. **E:** Testing boxplots of exploration time per video for each method as well as the human rater against each other: We found no significant difference from the human rater for all methods (paired Wilcoxon tests were used). **F:** Problem scheme when using larger and smaller objects and when resting on the object is excluded from data acquisition for object exploration. Left: Keypoint tracking software will appropriately not include data if the animal rests on a small object. Right: Keypoint tracking data may be falsely included if animals sit on large objects. **G:** Scatterplot showing the Spearman‘s correlations of exploration times measured by EXPLORE and the commercial software EthoVision and ANY-maze with human manual scoring. **H:** Heatmap revealing the Spearman‘s correlations between the assessed methods. **I:** Discrimination indexes (DIs) calculated for each of the methods: ANY-maze and EthoVision perform significantly different from EXPLORE and the human rater (paired Wilcoxon tests). **J:** No significant difference was found among all methods when comparing the detected exploration time per video (paired Wilcoxon tests were used. Asterisks showing the level of significanse: n.s. *p* ≥ 0.05, * *p* < 0.05).

### Losing tracking points by mirror images

Loss of tracking points due to mirror images can be avoided by a simple pre-processing step that includes cutting off the walls of the experimental arena in the video material before starting the analysis. However, this modification is non-trivial to apply without a pre-processing tool available. In contrast, EXPLORE does not require such pre-processing of data.

Results of EXPLORE correlated strongly to the human rater (*r* = 0.97, Spearman‘s correlation) in data with mirror images. Results of the keypoint tracking software EthoVision were less precise (*r* = 0.72, Spearman‘s correlation), as tracking point loss at reflecting walls of the experimental arena induced many outliers (Fig. 5B/C). This also compromised the calculation of the discrimination index (DI) in EthoVision, which was not significantly different from the 50%-chance level compared to all other methods (*p* < 0.05, Fig. 5D). We could further show, that tracking point loss due to mirror images significantly affected the exploration time: EXPLORE correctly measured the exploration time within the range of the human rater, while commercial software such as EthoVision and ANY-maze differed significantly (paired Wilcoxon tests were used, *p* < 0.05, Fig. 5E).

### Switching of head- and tail tracking-points when rodents have a similar colour to the object they explore

Our experiment shows that in contrast to EXPLORE keypoint tracking software such as ANY-maze and EthoVision have severe problems to process data where objects and mice have similar colours.

While results of ANY-maze and EthoVision poorly correlated to the human rater (*r* = 0.51, *r* = 0.5, Spearman‘s correlation), EXPLORE provided similar results compared to manual scoring by an expert (*r* = 0.92, Spearman‘s correlation) (Fig. 6B/C). Furthermore, DIs were significantly different for both keypoint tracking software compared to EXPLORE and the human rater (paired Wilcoxon tests were used, *p* < 0.05, Fig. 6D). No difference was found among all methods when quantifying the total exploration times per video (Fig. 6E).

### The choice of object size for exploration matters

If climbing or sitting on an object can be counted as exploration behaviour is still a matter of debate. However, if we assume to include these behavioural features, data with tracking points of the head will not account to the measured exploration time when e.g. the animal is sitting on a small object (∼ 2 cm): The head of the animal reaches beyond the contour of the object and data may thus not be included (Fig. 7A). In contrast, if we would like to exclude the behavioural feature ”sitting on the object” from object exploration, tracking the head of an animal sitting on a large object (∼ 10 cm) will fail to do so: Tracking points of the head will be within the contour of the object and data therefore not excluded (Fig. 7F). Furthermore, the calculation of the DI can be compromised if (two) objects provided in the arena differ in size, no matter if the researcher decides to in- or exclude behaviour with resting on objects.

To assess these issues we tested EXPLORE in comparison to keypoint tracking software (EthoVision, ANY-maze) and human rater accuracy if resting behaviour on the object was included in calculating object exploration (Fig. 7B-E). Results gained with ANY-maze and EthoVision (*r* = 0.86, *r* = 0.82, Spearman‘s correlation) correlated less to the human rater than EXPLORE (*r* = 0.95) (Fig. 7B/C). When calculating the DIs we found that results from ANY-maze and EthoVision significantly differed from the 50%-chance level (paired Wilcoxon tests were used, *p* < 0.05), while EXPLORE performed in the range of the human rater (Fig. 7D). The measured exploration time per video did not differ among methods used (Fig. 7E). We then excluded ”sitting on the object” as a behavioural feature of object exploration in our training data for EXPLORE (Fig. 7G-J). Results of EXPLORE also outperformed the results of common keypoint tracking (correlation to human rater for ANY-maze: *r* = 0.8, for EthoVision: *r* = 0.66 and EXPLORE: *r* = 0.95; Spearman‘s correlation)(Fig. 7G/H). The calculation of the DIs were again significantly different for EthoVision and ANY-maze compared to EXPLORE and the human rater (paired Wilcoxon tests were used, *p* < 0.05, Fig. 7I). We found no difference for the overall exploration time quantified by all methods (Fig. 7J).

### A set of GUIs for a step-by-step analysis of object recognition tests

EXPLORE provides users with all necessary tools for a successful object recognition test analysis, guided by three graphical user interfaces (GUIs): One GUI is designed to label training data and train the network (Training GUI). A second GUI can be run to perform the inference on all the experimental videos (trained and novel data, Prediction GUI) and a third GUI can be used to correct insufficient predictions by additionally labeling a few selected frames and re-feed the data to the network (Correction GUI).

In the Training GUI a user will first define project path and name, select videos to label from (randomly chosen or selected), define the number of frames to label, define names for behavioural classes along with corresponding keyboard keys to label frames with exploration behaviour. EXPLORE then shows the first frame to label. The user needs to draw a square around the surface of the arena, a square around each object (this will create the coloured boxes in the prediction videos later) or a small square somewhere in the image for behaviours (preferably in the corner where it is not overlapping an object). Then the user labels the frames where exploration behavior is identified. After that, EXPLORE automatically creates the training and validation data and the network will be trained.

After training the network, the Prediction GUI can be opened. A user can choose a project (of the trained data or another one) and the trained model file (.h5), define the outcome measure (in minutes) and select videos to run inference on. EXPLORE stores the results in a table as a .csv file and creates ”Prediction Videos” with coloured boxes around the objects or in the corner for behaviours when a frame was classified as such. This visualization helps the user to evaluate how well EXPLORE performed in the prediction step.

If the prediction was not satisfying enough, EXPLORE provides a GUI for correction. In the correction GUI users can define a project, the path to the raw videos, and the already processed ”Prediction Videos” created during the prediction step in the Prediction GUI. A user can then scroll through the videos and select a few sequences which the model falsely classified as exploration behaviour. These frames can be relabeled. EXPLORE will then incorporate the newly labeled data to the old data and re-train the network. After that, the prediction step can be performed again.

EXPLORE also provides a GUI to manually score video data. This might be helpful in particular in the beginning of the analysis so that users get experienced which kind of behaviour they would like to include or exclude as object exploration behaviour. With an additional GUI (see Supplementary Fig. 1) users can rapidly detect and analyse roaming behaviour in the arena: A very important preliminary step prior to the ORT is to assess if animals show preferences to some quadrants of the arena due to co-founders (e.g. local odours, differences in light conditions or temperature). If there is a preference, the outcome of an experiment can be biased and must thus be avoided. Therefore, researchers usually habituate animals to the experimental conditions and video tape roaming behaviour in the empty arena before presenting objects to them. The feature provided by EXPLORE takes mean pixel intensities of quadrants to calculate how long an animal explores each quadrant, allowing experimenters to easily quantify preference or avoidance of specific space within the arena and perform statistical tests (see Supplementary Fig. 1). The GUI allows to plot heat maps or line plots for individual animals (Supplementary Fig. 2) to immediately access and analyse if normal or divergent exploring behaviour can be found for individual animals: E.g. in an arena with two objects expected behaviour includes that animals spent more time in the quadrants with the objects than in those without (Supplementary Fig. 2).

## Discussion

We here present EXPLORE, a deep learning-based method, built into simple, user-friendly and open-source GUIs for the reliable analysis of object recognition tests in mice. EXPLORE achieves human expert-level accuracy in scoring exploration behaviour and was assessed on a number of different data sets. EXPLORE in particular out-performs expensive and commercially available software (EthoVision, ANY-maze) under specific conditions, where keypoint tracking software failed (Fig. 5, 6, 7). One of the great advantage of EXPLORE is that users can decide by themselves, which kind of behaviour to include or exclude as exploration behaviour by choosing and labeling the respective training data set, which is a prerequisite for reaching scoring results similar to human expert raters (Fig. 4). Furthermore, our software scores typical final outcome parameters such as the exploration time at defined objects to calculate the discrimination index, which is used by many neuroscience labs to quantify and compare cognitive performance. EXPLORE will thus save researchers great amounts of time, will lead to more reproducible results and will allow researchers to increase the number of experiments a lab can reasonably perform or the number of animals that can be investigated.

EXPLORE joins a growing community of open-source tools using deep learning to automatically quantify animal and human behaviour for biomedical research in a supervised or unsupervised manner [14, 15, 18, 21–24]. However, most of these tools have been designed for a general purpose of behaviour analysis and quantification, applicable to all kinds of videos with behaviour. In contrast, EXPLORE was specifically developed to provide an easy-to-use tool for the analysis of object exploration. Although it is in theory possible to label data and train our network to predict all kinds of behaviour in even different animal species - as the feature of training the network for specific behaviour already exists and has been tested here, e.g. ”stretching” or ”sitting on object” - we did not assess our tool on these further applications. EXPLORE acts as a stop-watch function of scoring specific (object exploration) behaviour, which is - at least to our knowledge - not present in any of the other tools but essential for the analysis of object recognition tests. These other tools are in general powerful but their adaptation to score object exploration requires at least some technical knowledge.

Moreover, we show that EXPLORE outperforms commercial software in many scenarios and reaches human accuracy. There are many commercial software packages on the market (i.e. ANY-maze, EthoVision, etc.). While they provide an enormous amount of possibilities to analyse a variety of experiments, they often lack the ability to capture complex ethological behaviour. Keypoint tracking software was not able to detect more complex behaviour e.g. ”sitting on the object” or ”stretching the body when being at the object”. In contrast, EXPLORE revealed a very robust and accurate performance close to the level of human experts (Fig. 4). Most of the keypoint tracking software make use of simple 2D-masks to detect when animals enter a specific zone around the object. As those software are unable to ”identify” specific exploration behaviour or do not allow the users to define ”exploration behaviour” by themselves and most often depend on geometric parameters in relation to the object, the pure presence of the animal within e.g. a certain radius around the object is accounted, potentially resulting in a false frame-count. In addition, large outliers are created and potentially falsify final outcome parameters such as the discrimination index (Fig. 5, 6, 7). While keypoint tracking software rely on high contrasts between arena, objects and animals in order to perform well, EXPLORE is not limited by any of these conditions (Fig. 2).

To counteract the kind of black-box behaviour of neural networks, we implemented several control mechanisms in the GUIs of EXPLORE to allow continuous monitoring of the training and prediction mode to detect exploration behaviour by the end-users including early stopping (to prevent over-fitting), plots of the training and validation metrics as a function of training epochs, and most importantly, videos with objects masked in different colours whenever a frame was predicted as object exploration behaviour. In case the trained network does not predict well on the remaining data set, users can correct the prediction with a few simple steps, re-train the network and re-predict on the data to obtain satisfying results (Fig. 2).

In summary, we present a simple, ready-to use, open source pipeline to perform the different analysis steps for object recognition tests with higher versatility, precision and comparable investment of time with expensive commercial software. EXPLORE robustly performs on different arena settings, background- or animal colours as well as on multiple amounts of objects used in the experiment. It is able to in- or exclude complex ethological behaviour from the analysis. We provide a set of GUIs ensuring accessability also for neuroscientists with no expertise in programming or deep learning. We here presented EXPLORE as a tool to primarily analyse object recognition, but it could potentially be used beyond this scope to quantify user-defined behavioural features, where a ”stop-watch function” for scoring behaviour is required.

## Materials and methods

### EXPLORE pipeline

Along with this publication we are releasing an open-source repository with code to manually label training data, train models, perform inference on new videos and correct inferences. The complete repository as well as corresponding user-friendly instructions for the GUIs can be found at https://github.com/victorjonathanibanez/EXPLORE.

### Implementation

We implemented EXPLORE in Python 3.6 [25]. We used the high-level machine learning API Keras [26] based on TensorFlow version 2.0 [27] for the architecture of our convolutional neural network, and Scikit- learn version 0.24.1 [28] for the evaluation metrics and the k-means algorithm. We used OpenCV (mac version: opencv-python 4.1.1.26, windows version: opencv 4.5.0) [29] for video or image reading and writing as well as for video or image cropping and resizing. Our graphical user interface was built with Tkinter, version 8.6.10 [30]. We tested our system on a Windows work station (Windows 10, NVIDIA GeForce GTX 1660 SUPER, 64 GB CPU RAM) and on a MacBook Pro (macOS catalina, 2.6 GHz Dual-Core Intel Core i5 processor, 8 GB RAM)

### Experimental set-up for data acquisition

#### Animals

Relevant variables of animal subjects, which should be carefully considered when performing an object recognition test (ORT), are strain, age, sex and estrous cycle [19]. For our experiments 9-12 months old adult B6/j-Rj mice (n=23) and white BALB/c mice (n=3) were obtained from Charles River, Germany. Mice were housed in standard cages (n=3-5 animals per cage) containing a sheet of paper tissue and a small red Plexiglas house for nesting behaviour. Animals were acclimatised to the housing room of the animal facility at least 1 week prior to the start of the experiments and had access to food and water ad libitum. The housing room was maintained at constant room temperature (22*±*1 °C) and under a reversed light-dark cycle (light 9pm - 9am). All experiments were carried out during the active cycle of the animals (dark period) and according to the guidelines of the Federal Veterinary Office of Switzerland and the license ZH 241 2018 approved by the Cantonal Veterinary Office in Zurich.

#### Object recognition test (ORT)

The object recognition tests were conducted in an acoustically isolated room in custom-made arenas (PVC fiber plates; white, or black squared: 40 × 40 × 40 cm and 50 × 50 × 50 cm / black round: radius = 20 cm) with custom made small objects built from little wooden pieces, screws, dowls, thumbtacks (2-4 cm), a larger black round wooden object (radius = 4 cm), a small wooden round (radius = 1.25 cm) and a large wooden round object (radius = 5 cm) as well as a larger squared object (4 × 4 × 10 cm). To assess any preference of the animals to explore specific parts of the arena or favor some objects over others, tests with *n* = 4 animals were performed prior to the actual object recognition test: Animals were placed for 10 min in the arena and roaming behaviour was video-taped by a camera (Panasonic HC-V180) at 25 frames/s. The camera was installed above the arena, ensuring a full overview of the whole arena. An LED strip (PHOBYA LED-Flexlight HighDensity) in the red-light spectrum surrounding the top of the arena provided illumination for the recordings in the setting of the squared white arena. The round black arena was illuminated with moderate room light. The squared black arena was illuminated from a transparent bottom with white light in an otherwise dark room. For the object recognition task, animals were acclimated to the experimental room and the empty arena for 2 min. at three consecutive days prior to the test (habituation phase). In the first part of the object recognition test - the acquisition phase - two identical objects were placed in two different quarters of the arena and animals were allowed to freely explore the arena and the objects for 4-6 minutes (depending on experiment). In the subsequent testing phase, one of the similar objects was replaced by a novel object and the animal was again allowed to explore the arena and objects for 4-6 minutes. The time between acquisition and testing phase (inter-trial period) was set to 24 h. For the evaluation of EXPLORE we did only consider one session, either acquisition- or testing phase.

#### Data and manual scoring

Our deep learning framework was tested on seven different data sets. Each data set was selected to assess different aspects of difficulty for the neural network to quantify exploration behaviour: Due to the different data sets we tested for diverse shapes of arenas, different background colours, different colours of mice, multiple sizes of objects, different amounts of objects as well as special behaviours such as ”stretching the body on the object” or ”sitting on object”. Each data set was manually scored by at least one expert with more than two years of experience to quantify exploration behaviour in an open field or object recognition task in mice.

#### Main experiment

The experiment to test the general performance of EXPLORE and to investigate misclassification was conducted in a white squared arena (see above in ”Experimental Setup”) with red light and small, wooden objects. In this experiment we used 15 different videos of black mice (female) with a length of six minutes each. For this main experiment, three experts manually scored the experiment twice to assess inter- and intra-variability between human raters. Before manual scoring, the raters discussed and agreed on how to score exploring behaviour: Exploring an object was counted as such when the nose of the animal was directed towards the object within a radius of 2 cm around the object. Climbing or being on the objects was not counted as exploration (see Fig. 1 and Supplementary Video 1).

#### Difficult light conditions: bright

To assess the correction function and performance of EXPLORE in difficult light conditions when mice and arena background are bright, a white squared arena, small wooden objects and white female mice (see above in ”Experimental Setup”) were used. In total, 12 videos with a length of 6 minutes each were analysed. All videos were recorded from the top view of the arena. All videos were manually scored by one expert rater. The scoring behaviour was classified in the same way as in the main experiment (see Fig. 2 and Supplementary Video 2).

#### Difficult light conditions: dark

The data to assess the correction function and performance of EXPLORE under dark light condition was videotaped from the top view, using a round black arena, small wooden objects and black female mice (see above in ”Experimental Setup”). Again, 12 videos with a length of 6 minutes each were analysed. The scoring behaviour was classified in the same way as in the main experiment. All videos were manually scored by one expert rater (see Fig. 2 and Supplementary Video 3).

#### Multiple objects / Mirroring errors

The data showing the performance of EXPLORE on multiple objects as well as highlighting the problem of mirroring errors in keypoint tracking software was coming from the same experiment: 8 videos of 4 minute length each, filmed top view in a squared black arena, with black male mice exploring 3 small wooden objects (see above in ”Experimental Setup”). This data was manually scored by an expert, in the same way as previously defined for the main experiment (see Fig. 3, Fig. 5 and Supplementary Video 4).

#### Complex behaviour

A set of 6 videos with a length of 5 minutes each was recorded. A white squared arena, red light, larger squared objects and black male mace were used and captured with a top-down view (see above in ”Experimental Setup”).. Here, for the manual scoring the same criteria were applied as in the main experiment, only ”complex behaviour” was scored differently: We defined ”stretching” as a behavioral feature when an animal elongates its body reaching with the head towards the top of the object until it is securely sitting on the object. We scored the time for ”sitting on object” until the mouse is at the verge of the object ready to jump from the object. The jump from the object was then again classified as ”stretching” as the animal has an elongated body until the save landing on the arena floor, where the body no longer is elongated. (see Fig. 4 and Supplementary Video 5).

#### Switching head- and tail tracking-points

To highlight the problem of switching head- and tail tracking-points when object and mice have similar colour, a white squared arena, black female mice and larger round, black objects were used (see above in ”Experimental Setup”). For this setting, 10 videos with a length of 6 min. each were filmed with a top-down view and the data was then manually scored as described in the main experiment by one expert (see Fig. 6 and Supplementary Video 6).

#### Large-/small object

The effects of large- and small object size in relation to the body length of the animal was investigated by using a white squared arena, black female mice, red light and different sized, wooden objects in separate video sessions (see above in ”Experimental Setup”). In total, 2 *×* 6 videos (1 single large object, 1 single small object) with a length of 5 minutes each were used. The videos were manually scored once by an expert as described in the main experiment (without animals on object) and once counting the animals being on the objects (see Fig. 7 and Supplementary Videos 7 and 8).

#### Data availability

Videos and human annotations are made available over the project website: https://github.com/victorjonathanibanez/EXPLORE

### Object exploration analysis with EXPLORE

A typical outcome parameter of the ORT is the time spent exploring an object in the arena during a fixed time period. Apart from manual scoring, automated approaches such as ANY-maze or EthoVision use keypoint tracking on the animal [11] to detect interactions with objects in the arena and quantify the exploration time. Although these approaches can be applied successfully, there are several disadvantages such as loss of tracking-points and the inability to distinguish between complex explorative behaviour (including ”sniffing” on the object) and pure roaming activity, resulting in time-consuming manual corrections. To overcome this hurdle, we created an easy-and-fast-to-use framework making use of supervised deep learning, with a small subset (e.g. 12000 frames / 8 minutes) of manually labelled frames as training data for the network.

#### General workflow

Our pipeline is designed to guide a user through the entire analysis process (see Fig. 8) with the help of graphical user interfaces (GUIs). After recording an experiment (Fig. 8 (1)), the first step is to choose all relevant videos that are part of the analysis and sample a subset of frames from the entire collection (Fig. 8 (2)). This step ensures that enough variability is captured for training subsequently. Users can decide between two options - either 1) random selecting or 2) users choose videos themselves with specific behaviour to be in- or excluded from the analysis (Fig. 8 (3)). For both options, a user-defined number of videos *n* with a length of *i* minutes and a user-defined target number of *j* minutes for the manual scoring video is created, where *i, j, n* ∈ ℕ, *n, j* ≥ 1 and *j < i × n*.

**Fig 8.**
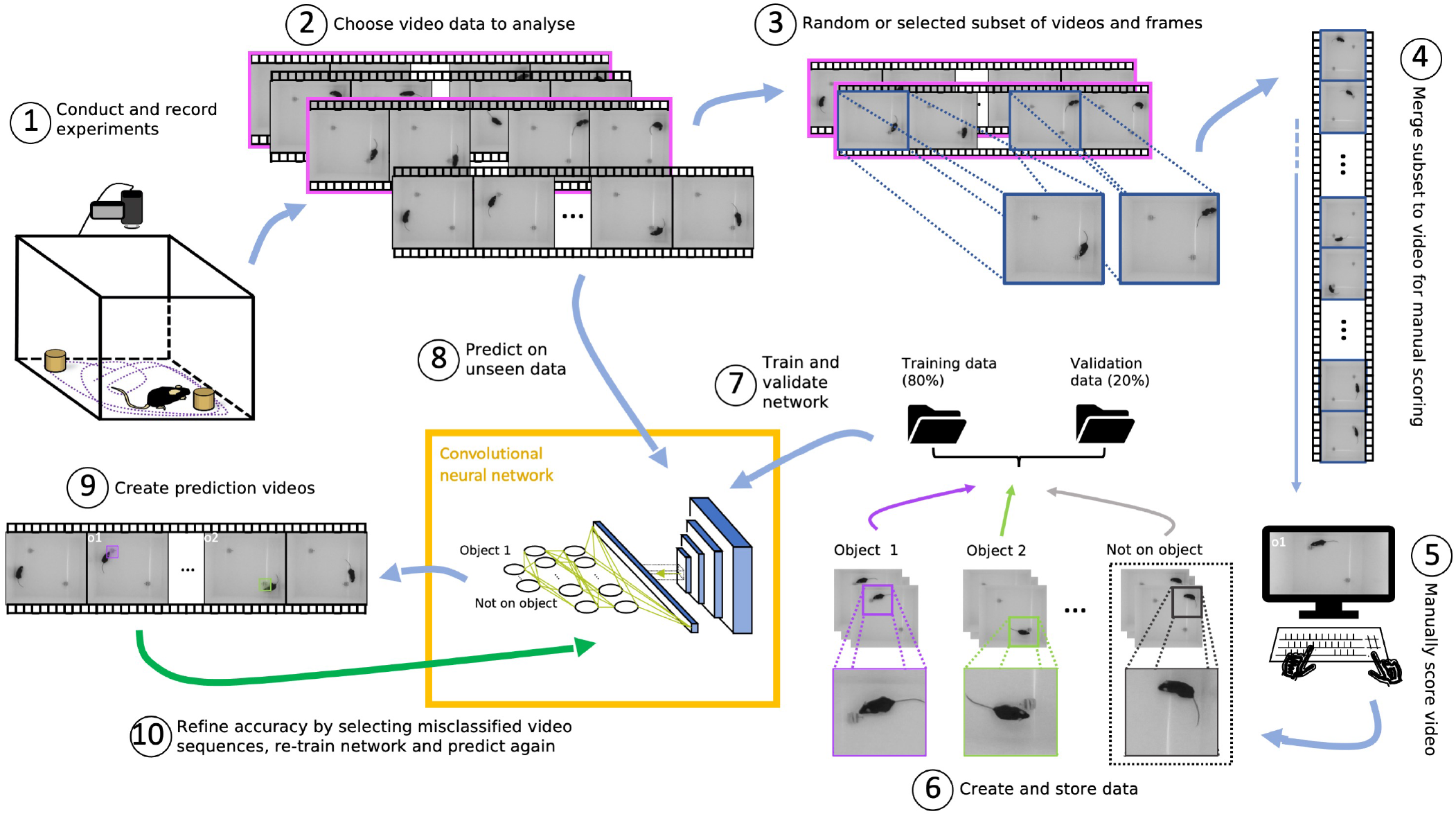
Overview of the EXPLORE pipeline. Deep learning framework for automated exploration analysis in object recognition tasks. **(1)** Experiments have to be recorded from the top view. **(2)** From the entire video data a subset is chosen. **(3)** From this subset, another subset of frames is selected **(4)** This final subset is then merged together to create a video for manual scoring. **(5)** The video is then manually scored. **(6)** When completed, the labeled data is split into training and validation data. **(7)** A convolutional neural network is trained on the training data and validated on the validation data. **(8)** The trained network can then be used to predict frames with exploration behaviour on the entire video data. **(9)** Videos are created with objects highlighted in different colours once the exploration is predicted at a given object in a frame. **(10)** If the prediction is not satisfying enough, the labeling can be corrected, the network re-trained and the prediction be repeated.

For 1) random selecting we implemented the k-means algorithm from Scikit-learn [28], which groups all videos into *n* clusters of related videos. From each cluster, one video will be randomly selected. If e.g. the camera is shifted slightly during the experiment, applying k-means is a good way to capture the resulted variability between the different videos.

2) Choosing videos manually allows to capture specific behaviour which only occur in a few videos of the experiment. From the selected *n* videos, a fraction *j/*(*i × n*) of each minute will be merged to the final manual scoring video. Thus, the temporal contiguity of the videos remains intact (larger fragments from videos instead of random frames) so that users can efficiently scroll through the videos for scoring.

For further setting-up the analysis, the user defines the labeling of the object(s) as well as a key on the keyboard which can be pressed during manual scoring for the training data when exploration behaviour is notified for the respective object. After drawing a rectangle around the arena surface (frames will be cropped to that size) and around each object (for the coloured mask in prediction videos), the manual scoring video will be merged and the user has to manually score exploration on the object by pressing the respective key (Fig.8 (4,5)). When the user finishes the manual scoring, the labeled frames will be resized to 150 *×* 150 pixels, split into training and validation data (80%, 20%) and the training will automatically start (Fig.8 (6,7)). When the training is finished, the trained network can be used to predict ”exploration behaviour” on the entire video data set and prediction videos will be created with coloured boxes around the objects in frames where exploration was predicted (Fig.8 (8,9)). This step allows users to validate how well the network predicted exploratory behaviour. In case there were inaccurate predictions, the sequences can be re-labeled and re-trained and finally re-predicted to achieve satisfying results in an iterative manner (Fig.8 (10)).

#### Model architecture and hyperparameters

In order to achieve best results and accelerate the analysis we specifically designed and tuned our custom convolutional neural network. The first building block of the model architecture consists of four convolution layers, each with a max pooling layer. The number of output filters doubles for each convolution layer from 32 to a final number of 256 output filters. For the convolution operation a kernel size of 3×3, followed by a max pooling operation with a kernel size of 2×2 was used. A flatten layer serves as transition from the first to the second building block. The fully connected building block consists of two dense layers with 500 nodes each, a dropout layer with a dropout rate of 0.5%, as well as an output layer representing the second building block. To introduce non-linearity, a rectified linear unit (ReLU) activation function was used for all layers, except the final one, where either a sigmoid (binary) or a softmax (multiclass) activation function was used to generate binary or multiple outputs (depending on the amount of objects to assess). For optimization, we use the adaptive moment estimation (Adam) algorithm [31] and the binary- or categorical cross-entropy loss function. We set the learning rate to 0.00015 and the batch size to 15, yielding the best results. The architecture is partially inspired by the architecture of the VGG16 network [32].

#### Training, validation and prediction

The network for each session was trained on 80% of the data and 20% of the data was used for validation. Rather than processing all data at once, data was loaded batch-wise into the random access memory (RAM) with the *flow from directory* function from Keras [26] for training and validating the network. Since there is most certainly always an imbalance between the classes in the data, weights were used for training and calculated as follows:

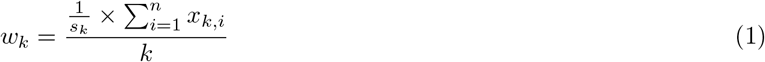

where *k* is the number of classes, *i* the *i*-th sample and *s*_*k*_ the number of samples for each class, with 1 ≤ *k* ≤ 6, *k* ∈ ℕ and *s, i* ≥ 0 and *s, i* ∈ ℝ. The built-in *accuracy score* from Keras [26] was used as a metric for validation. The network was set to train for a maximum of 50 epochs, with the *EarlyStopping* function from Keras [26] implemented to stop the training if the validation loss of the model reaches a minimum before the 50 epochs (patience = 5). Additionally, the *ReduceLROnPlateau* function from Keras [26] was used to reduce the learning rate by a given size if the validation loss does not further improve (patience = 3) to eventually converge to a lower minimum. After training the network on a subset of video frames, it can be used to predict on all of the experiment videos.

### Quadrant exploration analysis

Apart from the EXPLORE framework we also provide a tool to assess and quantify roaming behaviour in the arena (Supplementary Fig. 1). Before starting with the actual object recognition experiment, researchers usually want to investigate if animals show preference for a specific part of their experimental arena or if unexpected environmental co-founders occur which might influence the exploration behaviour. Our tool uses the mean pixel intensity for each quadrant of the arena. Other than preference, i.e. movement of animals throughout the arena (often measured as ”distance travelled”) can be quantified by investigating the frequency an animal transits from one quadrant to another of the arena. Abnormal behaviour such as freezing or preferences of specific local spots in the arena due to unwanted odor cues can be detected by analysing the time spent in each quadrant of the arena (Supplementary Fig. 2, for further details please see https://github.com/victorjonathanibanez/EXPLORE).

### Comparison to commercial software

The performance of EXPLORE was compared to commercially available software which are often used in neuroscience labs (ANY-maze by Stoelting Co.; Wood Dale, IL, USA and EthoVision by Noldus, Wageningen, the Netherlands [11]). Both software programs rely on tracking head-, mid- and tail-points from animals and drawing 2D-masks to mark an object zone. If a tracking-point (the head-point for object exploration) overlaps with a zone surrounding a given object *x* on a frame, this frame will be counted as ”exploration on object *x*”. For comparison to our manually scored training data for EXPLORE the same parameters were used for performance tests in ANY-maze and EthoVision (detection within a radius of 2 cm around the object).

### Statistics and evaluation

In order to evaluate the performance between EXPLORE, the two commercial software and the human raters (ground truth, Fig. 2) the time animals spent exploring the objects was calculated (a score in *s* per object, for each 1-*min* video segment) using the output (frames with detected exploration behaviour) of the different software and then compared with each other by computing the *Spearman‘s Correlation Coefficient (r)* which does not depend on linear assumptions and is less sensitive to outliers than other correlation measures. Other than that, the discrimination index (DI) was calculated as follows:

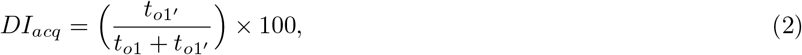

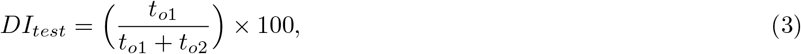

where *DI* stands for ”discrimination index” (with subscript *acq* or *test* for acquisition or testing session, respectively) in ORT, *t*_*o*1_ and *t*_*o*1_^*′*^ for the objects in the acquisition session and the equal object in the testing session and *t*_*o*2_ for the novel object in the testing session. The discrimination index was assessed (Fig. 2C) with one-sample t-tests (or Wilcoxon, if data was not normally distributed) against the 50%-chance level and a Bonferroni post-hoc test. Differences between the Spearman‘s correlation coefficient measures of EXPLORE, the commercial software and the raters were assessed with paired Wilcoxon tests. Differences in each class (e.g. ”object 1”, ”object 2”, etc.) between exploration time per video scored by the rater and exploration time per video inferred by EXPLORE were tested with paired Wilcoxon tests. To calculate differences between the exploration times animals spent in each quadrant of the arena (see EXPLORE quadrant analysis, Supplementary Figure 1) paired Wilcoxon tests and Bonferroni post-hoc tests were used. The level of significance was set at *p ≤ 0.05, **p ≤ 0.01, ***p ≤ 0.001.

## Supporting information

**Supplementary Fig. 1.**
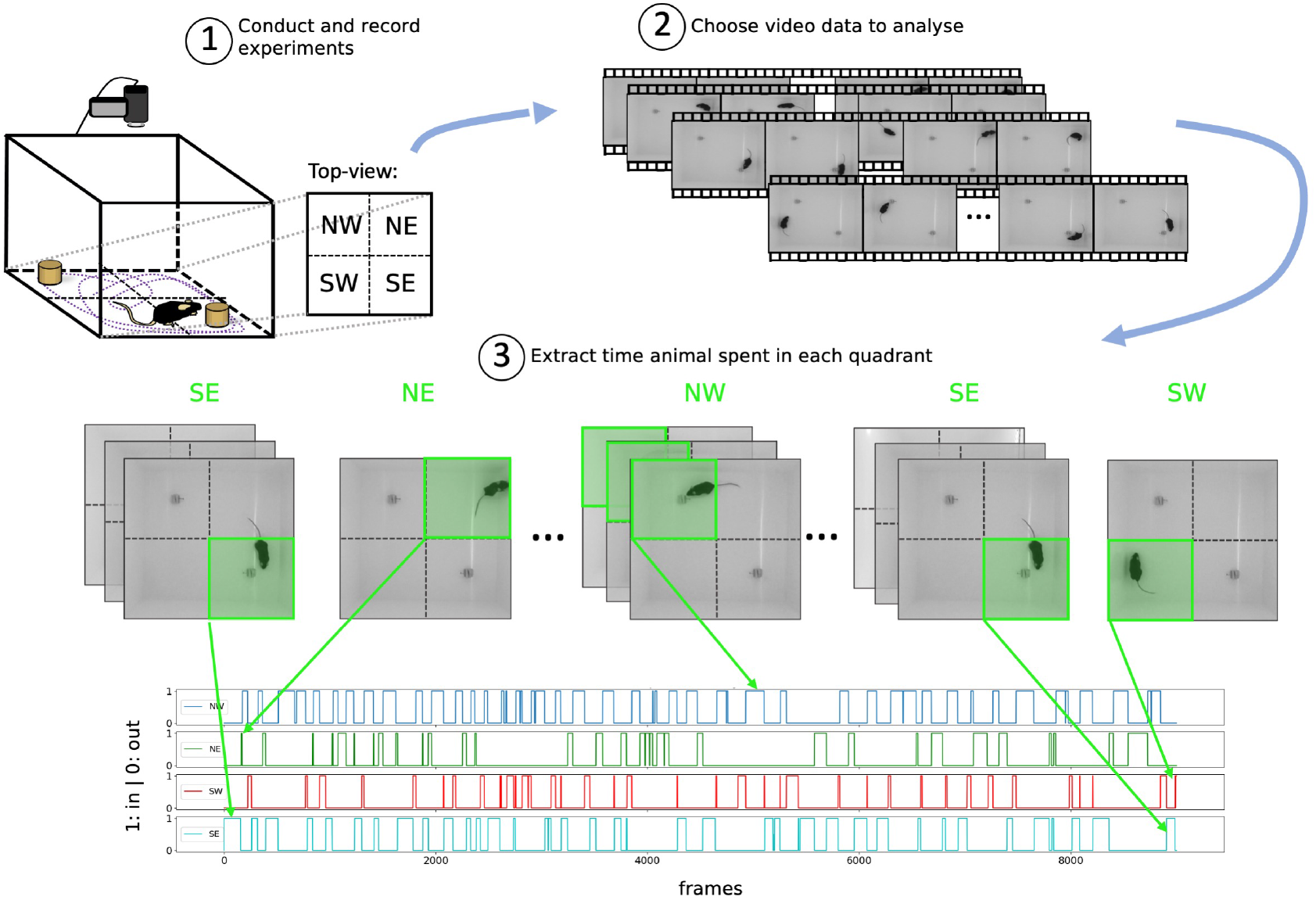
Overview of the quadrant exploration analysis. **(1)** Conduct and record experiments. **(2)** The videos to analyse can be selected. **(3)** In each frame, the mean pixel intensity for each quadrant is calculated. Then a threshold function is used to convert the values for each quadrant to ”0” or ”1” (the quadrant with the animal is assigned the value ”1”).

**Supplementary Fig. 2.**
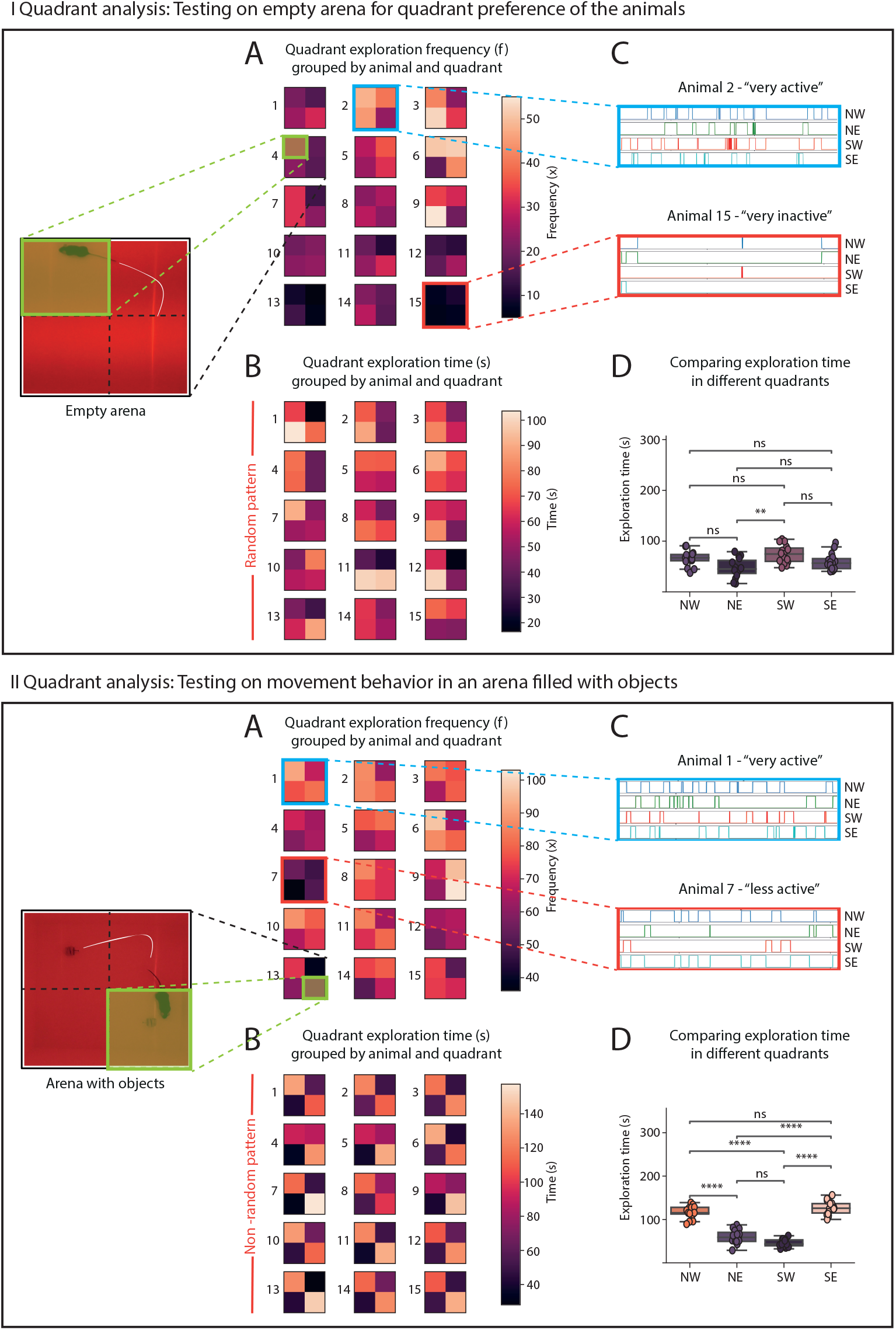
Quadrant exploration analysis can be used to qualitatively and quantitatively assess rodent behaviour and abundance in the quadrants of an experiment arena. **(I)** Running the quadrant exploration analysis on a set of experiment videos, where no object is in the arena. **A**,**B:** Heatmaps for the 15 assessed videos: Each numbered square represents the summed exploration frequency (**A**) or the summed exploration time (**B**) per animal with the four quadrants of the experiment arena (north-west (NW), north-east (NE), south-east (SE), south-west (SW)). **C:** two examples (2 animals) of the frequency plots for each of the quadrants: Animal 2 shows a lot of activity overall, animal 15 not much. **D:** To assess if the animals spent equal time in each quadrant and had no preference, the total exploration time for each quadrant can be compared: We used a paired Wilcoxon test with bonferroni post-hoc and found no significant difference between the quadrants. In **B** it can also qualitatively be seen that there is no ”pattern” visible: Animals seem to have randomly visited the quadrants. **(II)** As a placeholder for general cues that could potentially bias the experiment, we run the quadrant exploration analysis on a set of experiment videos, where 2 objects are located NW and SE in the arena. **A**,**B**,**C:** same as in **I. D:** Here, it can be seen that the summed exploration times for each quadrants are significantly different between NW and NE, NW and SW, NE and SE, SW and SE. **B** also shows that there is a clear ”pattern” visible: Animals explored NW and SE (object locations) longer than NE and SW. Asterisks showing significance level: ns ≥ 0.05, * ≤ 0.05, ** ≤ 0.01, *** ≤ 0.001, **** ≤ 0.0001.

**Supplementary Video 1. *Example video of EXPLOREs performance on our main data set***. *EXPLORE correctly predicts when the animal is exploring the two objects (Whenever EXPLORE counts exploration behavior, the objects are marked with a coloured frame)*.

**Supplementary Video 2. *EXPLORE video example on data set ”difficult light conditions: bright”***. *While keypoint tracking software fails, EXPLORE is able to classify the behaviours as exploration (Whenever EXPLORE counts exploration behavior, the objects are marked with a coloured frame)*.

**Supplementary Video 3. *EXPLORE video example on data set ”difficult light conditions: dark”***. *As in Supplementary video 2, EXPLORE sufficiently detects exploration behavior, even under these difficult light conditions (Whenever EXPLORE counts exploration behavior, the objects are marked with a coloured frame)*.

**Supplementary Video 4. *Sample video of EXPLORE performing on multiple objects*** *EXPLORE is able to correctly classify behaviour on multiple objects. Whenever EXPLORE counts exploration behavior, the objects are marked with a coloured frame*.

**Supplementary Video 5. *A video sample revealing a more versatile usage of EXPLORE***. *EXPLORE is able to precisely capture different complex features of behaviour, defined by the user (Whenever EXPLORE counts exploration behavior, the objects are marked with a coloured frame, behaviour are marked with small coloured frames)*.

**Supplementary Video 6. *Video sample demonstrating the performance of EXPLORE, where keypoint tracking software would lose or switch tracking points***. *EXPLORE correctly predicts when the animal is exploring the two objects (Whenever EXPLORE counts exploration behavior, the objects are marked with a coloured frame)*.

**Supplementary Video 7. *Video sample depicting the prediction of exploration behavior by EXPLORE when using small objects***. *EXPLORE captures what the user defines as exploration behavior independent of the size of the object, leading to correct labels (Whenever EXPLORE counts exploration behavior, the object is marked with a coloured frame)*.

**Supplementary Video 8. *Video sample depicting the prediction of exploration behavior by EXPLORE when using large objects***. *EXPLORE captures what the user defines as exploration behavior independent of the size of the object, leading to correct labels (Whenever EXPLORE counts exploration behavior, the object is marked with a coloured frame)*.

## Acknowledgments

The authors thank Peter Rupprecht for advice in writing and structuring the manuscript and data.

